# Cholesterol Remodeling by CH25h Rewires IFITM3 Trafficking and Secretion Without Enhancing Antiviral Restriction

**DOI:** 10.64898/2025.12.09.693135

**Authors:** Yuxin Song, Julien Burlaud-Gaillard, Philippe Roingeard, Andrea Cimarelli

**Author notes:** Correspondent footnote: A. Cimarelli, Centre International de Recherche en Infectiologie, 46 Allée d’Italie, Lyon 69007, France.

## Abstract

Type I interferon induces multiple protein effectors to inhibit viral replication. Among these, the interferon-induced transmembrane protein (IFITM3) and the cholesterol-25-hydroxylase (CH25h) converge on the inhibition of viral entry by altering the behavior of membranes. Here, we dissect the functional and mechanistic relationship between these two membrane-acting effectors using HIV-1 and vesicular stomatitis virus (VSV) as models. We show that IFITM3 and CH25h restrict viral entry with similar efficiency, but act in a largely redundant manner during infection. Their redundancy is consistent across infection systems, cell types, and entry assays, indicating that both factors converge on a shared biophysical block to membrane fusion. Unexpectedly, we uncover a second layer of interplay in which the CH25h-25HC axis remodels IFITM3 trafficking. Exposure to 25-hydroxycholesterol drives IFITM3 from endolysosomal compartments to the plasma membrane, impairs its internalization into early endosomes, and increases its secretion in exosomal vesicles. These effects occur independently of direct antiviral activity and reveal that CH25h regulates IFITM3 cellular dynamics. Together, our study identifies a previously unrecognized cross-regulatory circuit between two IFN-induced antiviral pathways, highlighting how lipid remodeling by CH25h, and potentially by other cellular factors, can contribute to control the behavior of IFITM3.

**Importance:** Type I interferons induce numerous antiviral factors that frequently target the same vulnerable steps of viral infection, but how these factors influence one another is not well understood. IFITM3 and the CH25h are two potent, broad-acting inhibitors of viral membrane fusion. Here, we show that although their antiviral activities are largely redundant against HIV-1 and VSV, CH25h profoundly alters the biology of IFITM3. The product of the enzymatic activity of CH25h, the oxysterol 25HC, redistributes IFITM3 from endosomal compartments to the plasma membrane and enhances its release in exosomal vesicles— effects independent of direct antiviral activity. These findings reveal an unexpected layer of cross-regulation between lipid-modifying enzymes and membrane-embedded restriction factors. More broadly, they highlight how interferon-driven lipid remodeling can reshape the trafficking and secretion of antiviral proteins, expanding the functional landscape of innate immunity beyond viral entry inhibition.

## Introduction

Type I interferon (IFN-I) responses constitute the first line of defense that cells deploy against viral infection. Once triggered, hundreds of IFN-induced gene products then interfere either directly, or indirectly, with viral replication. Among these, two antiviral effectors stand out for their capacity to restrict viral entry by altering the properties of membranes: the interferon-induced transmembrane proteins (IFITMs) and the cholesterol-25-hydroxylase (CH25h).

IFITM1, IFITM2, and IFITM3 are the most prominent antiviral members of the IFITM family. IFITMs are membrane-bound proteins and share a typical domain organization with one intra– and one trans-membrane domain separated by an intracellular loop (IMD, TMD and CIL, respectively). They also possess N and C-termini of variable lengths that tightly regulate IFITM trafficking, stability, and post-translational modifications (1–3).

Functionally, IFITMs inhibit viral replication at two distinct stages. In target cells, endosome-resident IFITMs block low-pH dependent fusion between viral and endosomal membranes, leading to virion degradation (4–8). In virus-producing cells, IFITM expression results in the release of virions that are intrinsically less fusogenic and therefore less infectious (9–11). Although the precise mechanism underlying this fusion defect remains incompletely understood, current evidence suggests a combination of direct membrane-rigidifying effects driven by IFITM insertion and multimerization, coupled with possible indirect effects on membrane composition (5–7, 12–14).

CH25h represents another IFN-induced antiviral defense acting at membranes. This endoplasmic reticulum (ER)–resident enzyme catalyzes the conversion of cholesterol into 25-hydroxycholesterol (25HC), a soluble oxysterol with paracrine and autocrine immunomodulatory functions (15–17). 25HC is a potent broad-spectrum antiviral factor and, similar to IFITMs, inhibits viral entry by impairing virus-to-cell membrane fusion—an effect thought to arise from the incorporation of its bulky, hydroxylated head group into cellular membranes (16, 18, 19).

Because IFITMs and CH25h inhibit a wide range of viruses at the same early fusion step, we sought to determine whether they act redundantly, synergistically, or influence each other’s cellular behavior. The results we have gathered indicate that IFITM3 and CH25h exert largely redundant effects on the replication of representative members of the *Retroviridae* and *Rhabdoviridae* (HIV-1 and VSV), under the conditions used here.

However, we identify a previously unrecognized connection between the two pathways: the CH25h– 25HC axis substantially redistributes IFITM3 from internal endosomal compartments to the plasma membrane. This relocalization correlates with an increased secretion of IFITM3 in exosomal vesicles. These findings reveal a novel mode of cross-regulation in which CH25h influences the trafficking and extracellular release of IFITM3—independently of their overlapping antiviral activities.

Overall, our study demonstrates that although IFITM3 and CH25h do not act additively to restrict HIV-1 or VSV infection, they influence each other’s biological properties in ways that may have broader implications for membrane biology, innate immunity, and vesicle trafficking.

## Results

### CH25h and IFITM3 restrict HIV-1 in a largely non-cooperative manner

To define how CH25h and IFITM3 functionally relate during the HIV-1 life cycle, we first examined viral spread using replication-competent NL4-3 virus in SupT1 cells expressing doxycycline-inducible IFITM3 and treated with concentrations of 25HC ranging from 0.17 to 1.5 μM (higher concentrations were avoided following pilot experiments pointing to toxicity in this cell type).

25HC treatment did not affect the steady state accumulation of IFITM3 as assessed by WB, nor altered cell proliferation in the timeframe of our spreading infection assays (Appendix Figures A1A and A1B). In this long-term replication setting, both IFITM3 expression and 25HC treatment reduced HIV-1 spread and in the case of 25HC, inhibition was proportional to its concentration. When combined, 25HC treatment of IFITM3-positive cells resulted in a stronger inhibition than either alone, but this occurred only at the highest concentration of 25HC used (Figure 1A). This effect could be due to either a true synergistic effect between IFITM3 and 25HC, or else to pleiotropic effects of 25HC. To discriminate among these possibilities, we measured their effects on the early and late stages of infection using single-cycle assays, according to the scheme of Figure 1B. To examine the early phases of the viral life cycle, HEK293T-CD4-CXCR4 cells expressing IFITM3 and/or a wild-type mCherry-CH25h protein alone or in combination were challenged with single-round of infection-competent HIV-1 viruses pseudotyped with the HIV-1 Envelope protein derived from the NL4-3 strain and coding for a GFP reporter gene. A catalytically inactive CH25h mutant (CH25h-M, containing the following mutations: H242E/H243E/H246E/H247E) was included as control (20). Two-three days post infection, cells were either analyzed by WB (Figure 1C), or by flow cytometry to determine the extent of infection (Figure 1D). Under these conditions, each protein was robustly expressed and no detectable changes were observed when the two proteins were expressed together (Figure 1C). As expected, expression of either IFITM3 or CH25h significantly inhibited the early steps of HIV-1 infection (Figure 1D). CH25h expression led to higher inhibition of HIV than IFITM3 (18.0% versus 57,3% respectively, Fig 1D for normalized values and Appendix Figure A1C for non-normalized ones), while the catalytically inactive CH25h-M mutant displayed no significant antiviral activity. Co-expression of IFITM3 and CH25h did not further enhance inhibition beyond the activity of CH25h alone (37.9%), and if anything, it was lower when compared to CH25h alone (18.0%), albeit this difference did not reach statistical significance.

**Figure 1.**
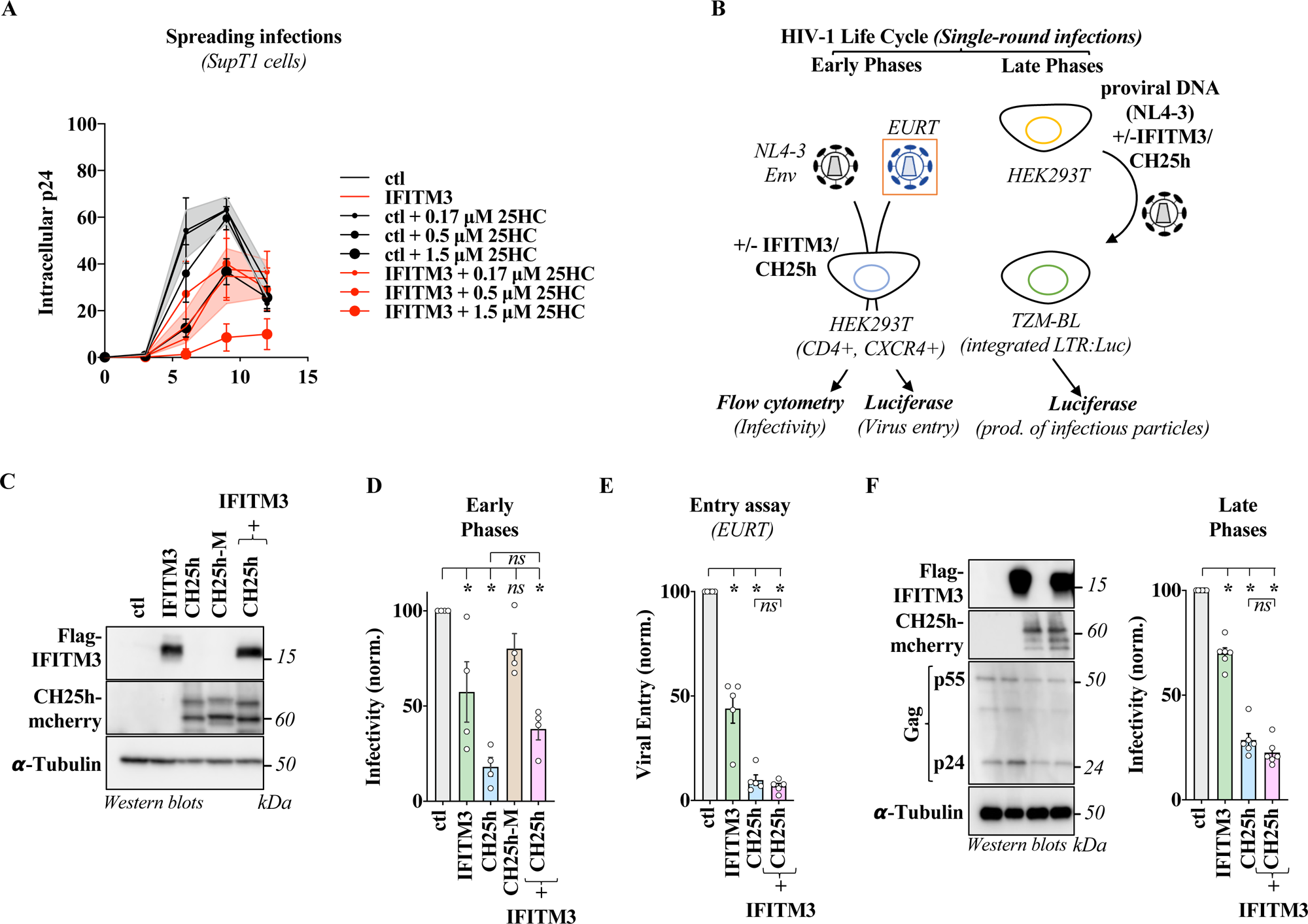
IFITM3 and CH25h exert largely redundant roles during HIV-1 infection. A) Control or dox. inducible IFITM3 stable SupT1 cells were treated with doxycycline (1 μg/mL), +/− 25HC prior to viral challenged with an MOI of 0.1. Cell aliquots were harvested at the indicated time points and the percentage of infected cells was measured by FACS following intracellular p24 staining. B) Experimental scheme used here to examine the role of IFITM3 and CH25h on the early and late phases of HIV infection. In the former, HEK293T expressing the CD4/CXCR4 HIV-1 receptor/co-receptors were first transfected with plasmids coding the indicated combinations of Flag-tagged IFITM3, *wild-type* CH25h-mCherry or its catalytically-inactive mutant CH25h-M-mCherry. Thirty-six hours later, cells expressing the different proteins were either analyzed by WB (C), or challenged with a single-round of infection-competent NL4-3 Env-pseudotyped-HIV-1 virus coding a CMV:GFP reporter cassette. The extent of infection was determined two-three days later by flow cytometry (D). To more specifically measure defects in virus entry, viruses incorporating a viral genome mimic coding for the Firefly Luciferase that can be directly translated upon entry of the virus into the cell were used to challenge target cells for eighteen hours in the presence of the capsid destabilizing agent PF74 (10 μg/mL). The extent of virus entry into the cells was measured in function of the accumulation of Luciferase activity in cell lysates (E). The late phases of HIV-1 infection were studied by directly producing HIV-1 viruses in HEK293T cells by coexpression of the DNAs indicated above together with the proviral DNA construct NL4-3 which allows the production of fully replicative virus. Infectious virus was harvested from these producing cells, diluted 100 times to nullify effects of 25HC on the new cycle of infection and used to challenge TZM-BL target cells. The extent of production of infectious virus was determined after measurement of the luciferase activity thanks to the LTR:Luciferase reporter integrated in TZM-BL cells (F). The WB panels depict typical results obtained, while graphs present AVG and SEM from independent experiments (C, n=4; D, n=4; E, n=5; F, n=6). * and ns, p<0.05 or non-significant following Ordinary one-way Anova, Dunnett’s multiple comparison tests.

To confirm that viral inhibition mapped at the step of HIV-1 entry into the cell, we used the entry/uncoating EURT assay, in which the translation of a virion-incorporated luciferase minigenome serves as a direct marker of virus entry into the cell (21). Cells are challenged with this virus and capsids are destabilized by incubating target cells with PF74, prior to Luciferase quantification eighteen hours later. Under these conditions, we found that IFITM3 and CH25h reduced early HIV-1 entry (Figure 1E), in line with data present in the literature (4–8, 16). However no cumulative effects were observed upon co-expression of the two.

Next, we assessed effects on late stages of replication by producing virions from cells expressing IFITM3 and/or CH25h and measuring infectivity of newly-produced virion particles on TZM-BL reporter cells (scheme of Figure 1B and Figure 1F). After forty-eight hours, the virus was harvested, diluted 100 times and then used to challenge TZM-BL target cells that bear an integrated LTR:luciferase reporter. The dilution step was introduced following pilot experiments to discard effects of soluble 25HC present in the supernatant of virus-producing cells that could have influenced the interpretation of the results by influencing virus entry in target cells, rather than the infectivity of newly-produced virions. Virus infectivity was measured twenty-four hours after viral challenge of TZM-BL cells, by quantifying the amount of luciferase produced which in these cells is directly linked to the production of Tat from newly-integrated proviruses. Under these conditions, virions produced in the presence of IFITM3 or CH25h exhibited reduced infectivity (66.0% and 24.2% of control, respectively). Again, co-expression resulted in inhibition comparable to CH25h alone (18.9%), indicating limited cooperation at this stage.

Together, these findings reveal first an undescribed effect of the CH25h-25HC axis on the late phases of HIV-1 and second, they demonstrate that IFITM3 and CH25h restrict HIV-1 through overlapping mechanisms, with negligible additive effects during single-cycle infection. Therefore, the additive effects observed during spreading infections are likely a consequence of the pleiotropic effects that 25HC may have on the cell physiology upon prolonged culture.

### CH25h and IFITM3 inhibit VSV infection in a similarly redundant fashion

Because IFITM3 and CH25h target membrane fusion, we assessed whether their relationship was conserved for viruses with different entry and replication strategies. To this end, we used a replicative VSV (Indiana serotype) bearing a GFP reporter transcription unit inserted between M and G (22). HEK293T cells expressing IFITM3, CH25h (WT or M) alone or in combination were challenged with a fixed amount of VSV and the extent of infection was then measured twenty-four hours later by flow cytometry (Figure 2A for WB and normalized infection data and Appendix Figure A2A for non-normalized infectivity data). As observed for HIV-1, both IFITM3 and CH25h markedly decreased susceptibility to VSV (Figure 2A, 61.1% and 35.9% of control, respectively), whereas mutations altering the enzymatic activities of CH25h led to a loss of antiviral activity (90.99% of control). Co-expression of IFITM3 and CH25h inhibited infection but did not further increase restriction compared with CH25H alone (28.07%). These effects were maintained across different viral inputs (Figure 2B) and were not cell type dependent, as similar effects were also obtained in A549 cells (Appendix Figure A2B).

**Figure 2.**
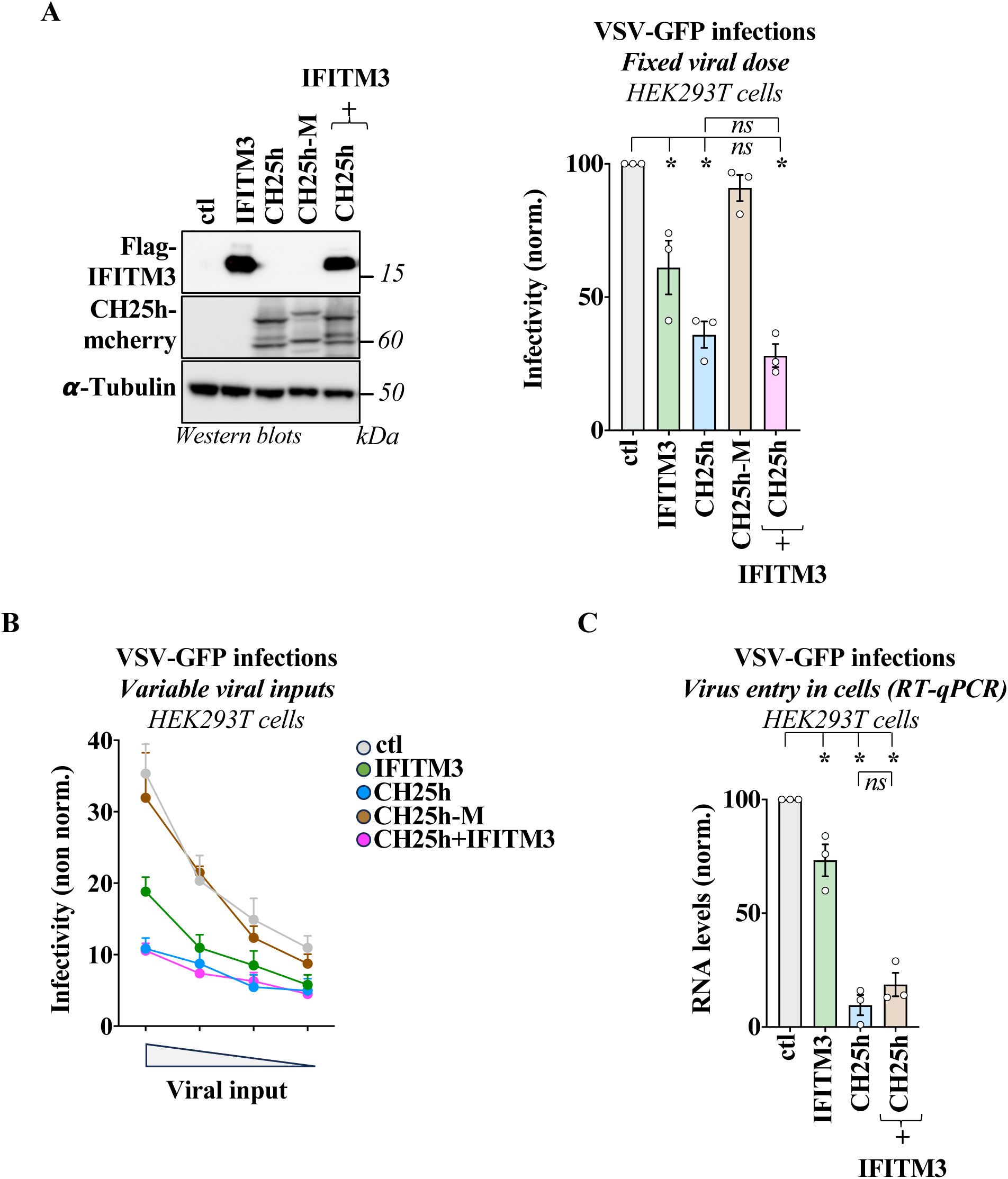
IFITM3 and CH25h exert largely redundant roles also against VSV. A) HEK293T cells transfected with the indicated plasmids were either collected for WB analysis or challenged with VSV-GFP at an MOI 0.05 for 16 hours, followed by flow cytometry analysis. B) A similar setup was employed using dilutions of the viral input. C) The extent of virus entry into the cell was determined by measuring the levels of VSV-GFP RNA in cells twenty min after viral challenge. Cells were trypsinized and washed to remove unbound virus. The WB panels presents typical results obtained. The graphs present AVG and SEM from independent experiments (A and C, n=3; B, n=6). * and ns, p<0.05 or non-significant, respectively, following Ordinary one-way Anova, Dunnett’s multiple comparisons tests.

To map the inhibition step, we quantified incoming viral RNA twenty minutes post-infection in the presence of cycloheximide to block secondary RNA synthesis (Figure 2C). Both IFITM3 and CH25h reduced the accumulation of incoming nucleocapsids, confirming a block at viral entry. Together, these results show that IFITM3 and CH25h restrict VSV entry in a largely non-additive manner, consistent with a shared mechanism of fusion inhibition.

### IFITM3 does not modulate the secretion of 25-hydroxycholesterol

To determine whether IFITM3 alters CH25h enzymatic output, we analyzed the antiviral activity of conditioned supernatants from cells expressing IFITM3, CH25h, CH25h-M, or both proteins. To this end, we first performed a dose response curve to soluble 25HC in HEK293T cells and assessed effects on both cell toxicity and susceptibility to infection (Appendix Figures A3A, A3B and A3C). Under our experimental conditions, we determined that cell toxicity was apparent only at concentrations of 25HC above 6.25 μM in this cell type (Appendix Figures A3A, A3B), while the susceptibility to infection of target cells incubated with different concentrations of soluble 25HC was inversely proportional to the concentration of 25HC and reached a plateau already at 0.195 μM (Appendix Figures A3C).

Next, to evaluate whether IFITM3 affected the levels of 25HC produced or secreted from cells expressing CH25h, the supernatant of cells was harvested twenty-four hours after transfection, filtered to remove cellular debris and directly used (with no dilution) to incubate non-transfected HEK293T cells. After additional twenty-four hours, the cells were challenged with VSV-GFP and then analyzed by FACS to examine their susceptibility to infection. In this assay therefore, 25HC secretion is measured indirectly through its effects on virion infectivity (Figure 3A). Under these conditions, supernatants from CH25h-expressing cells—but not from CH25h-M or IFITM3 alone—conferred strong resistance to VSV when applied to naïve cells, indicating the presence of biologically active secreted 25HC (Figure 3B). Importantly, supernatants from cells co-expressing IFITM3 and CH25h displayed antiviral activity indistinguishable from CH25h alone, demonstrating that IFITM3 does not influence 25HC production by CH25h, or its secretion.

**Figure 3.**
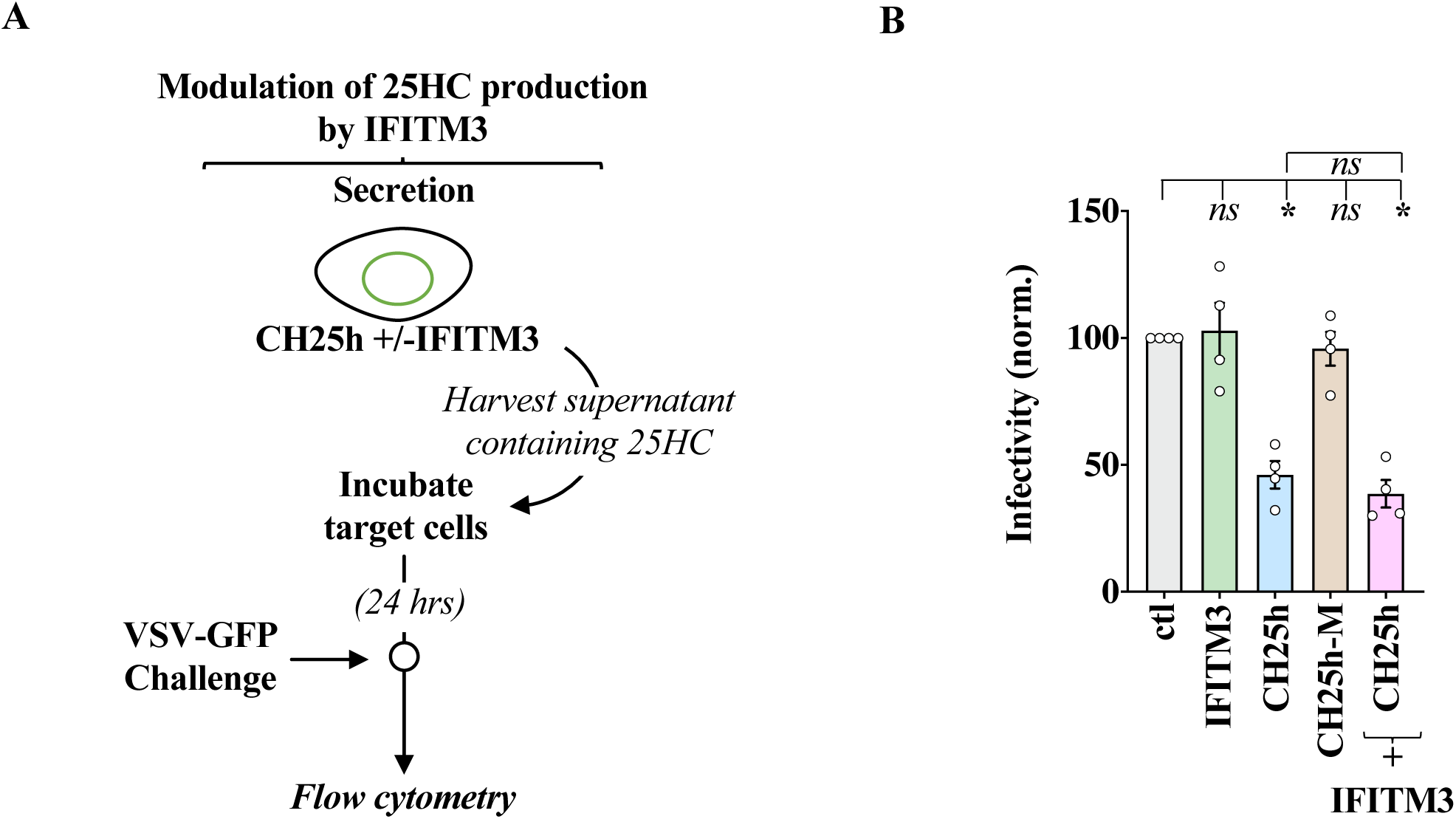
IFITM3 does not modify the extent of 25-Hydroxycholesterol production and release from CH25h-producing cells. A) Experimental scheme used here. B) The supernatant of HEK293T cells expressing the indicated proteins was collected twenty-four hours after transfection and used undiluted on untransfected cells for additional twenty-four hours. Cells were then challenged with VSV-GFP prior to flow cytometry analysis eighteen hours afterwards. The graph presents AVG and SEM of 4 independent experiments. * and ns, p<0.05 or non-significant, respectively, following Ordinary one-way Anova, Tukey’s multiple comparisons tests between the indicated conditions.

Despite the fact that we have not directly measured the levels of 25HC by mass spectroscopy, the observation that supernatants derived from cells expressing the catalytically-inactive form of CH25h fail to induce an antiviral behavior in target cell is a strong argument indicating that the antiviral effects measured in this context are essentially mediated by this oxysterol.

### The CH25h-25HC axis redistributes IFITM3 from endo-lysosomal compartments to the plasma membrane

We next asked whether CH25h and 25HC could instead influence IFITM3 behavior and to address this issue we examined the intracellular distribution of IFITM3 in cells exposed to high concentrations of 25HC (1.5 μM). Under these conditions, we observed a shift of IFITM3 from its canonical endosomal/vesicular pattern to the plasma membrane. Co-staining of cells with FITC-wheat germ agglutinin conjugates (WGA-FITC) confirmed a significant increase in plasma membrane–associated IFITM3 across multiple cell types (HEK293T, A549, SupT1, Figure 4A for representative pictures and 4B for quantitative analyses).

**Figure 4.**
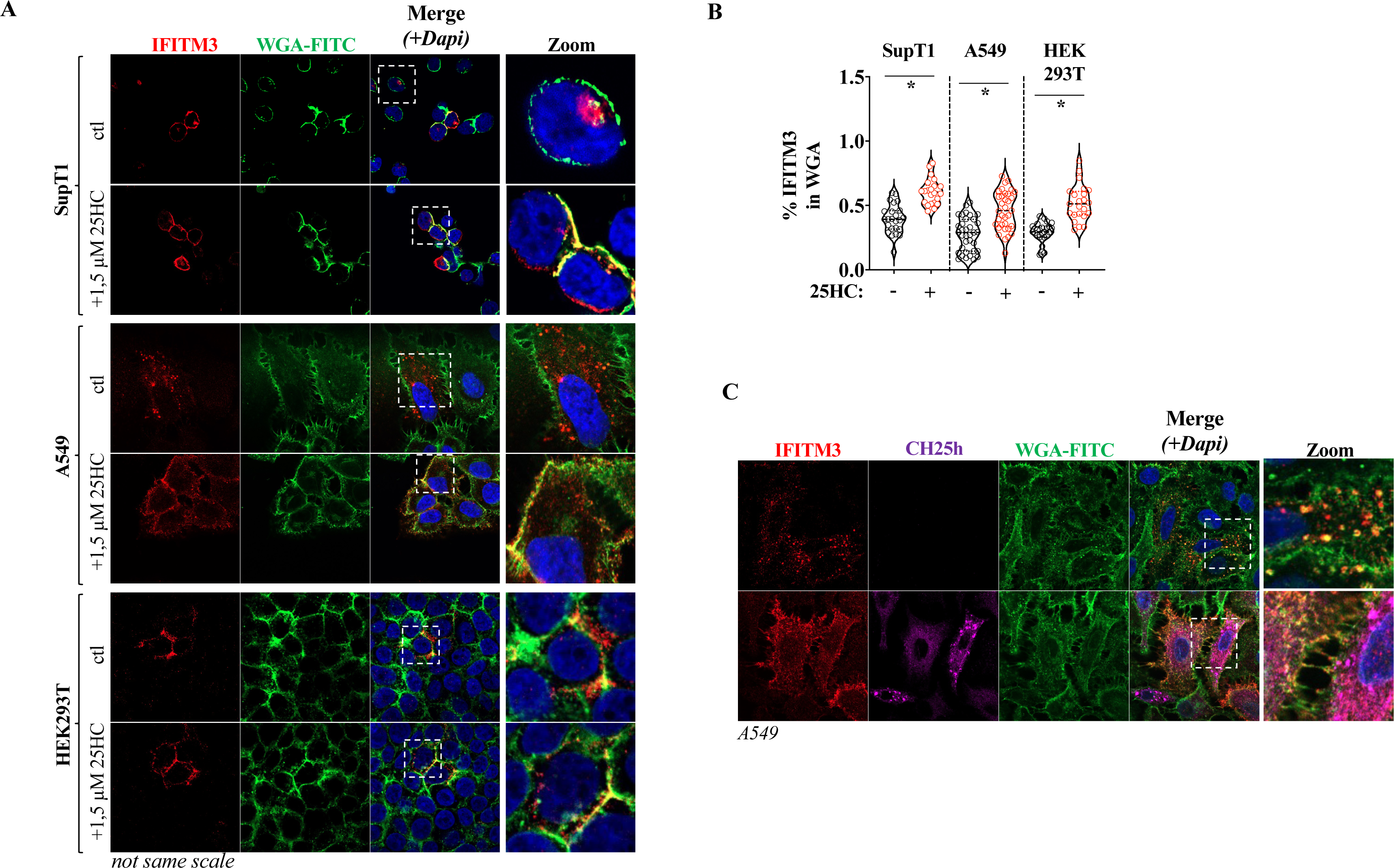
The CH25h-25HC axis promotes a redistribution of IFITM3 to the plasma membrane in several cell types. A) Expression of IFITM3 was induced upon incubation with doxycycline in cell pools of SupT1, A549 and HEK293T obtained following retroviral-mediated gene transduction. Cells were further incubated or not with 25HC (1.5 μM) for twenty-four hours prior to a 20 min labeling step at 37°C with FITC-WGA and confocal microscopy analysis. B) The graph presents the quantification of the percentage of the IFITM3 signal in the FITC-WGA one (Mander’s) obtained in 4 to 5 independent experiments and in 22 to 35 individual cells per conditions. *, p<0.05, according to a two-tailed, unpaired Student t test between the indicated conditions. C) A549 cells stably expressing dox.-inducible forms of IFITM3 alone or together with CH25h were induced with doxycycline (3 μg/mL) for twenty-four hours. Cells were then labeled with FITC-WGA as above, prior to confocal microscopy analysis.

To reinforce this observation, we examined A549 cells stably co-expressing both IFITM3 and CH25h. As with exogenous 25HC treatment, CH25h expression induced a robust redistribution of IFITM3 toward the cell surface (Figure 4C). Overall, these findings indicate that the 25HC-CH25h axis regulates the intracellular trafficking of IFITM3, independently from viral infection.

### 25HC reduces internalization of IFITM3 into early endosomes

Upon expression, IFITM3 is thought to reach the plasma membrane, where it is then endocytosed into early and then late endosomal vesicles thanks to specific sequences present in the long N-terminus of IFITM3 (23, 24). We therefore hypothesized that this step could be compromised by the 25HC-CH25h axis. To determine whether this was the case, we determined the amount of IFITM3 in early endosomes following confocal microscopy analysis in A549 cells exposed or not to 25HC. While when expressed alone, IFITM3 exhibited a consistent localization with the early endosomal marker EE1A, this colocalization was essentially lost in the presence of 25HC (Figure 5), therefore indicating that 25HC induces an early defect in the recycling of IFITM3 from the plasma membrane.

**Figure 5.**
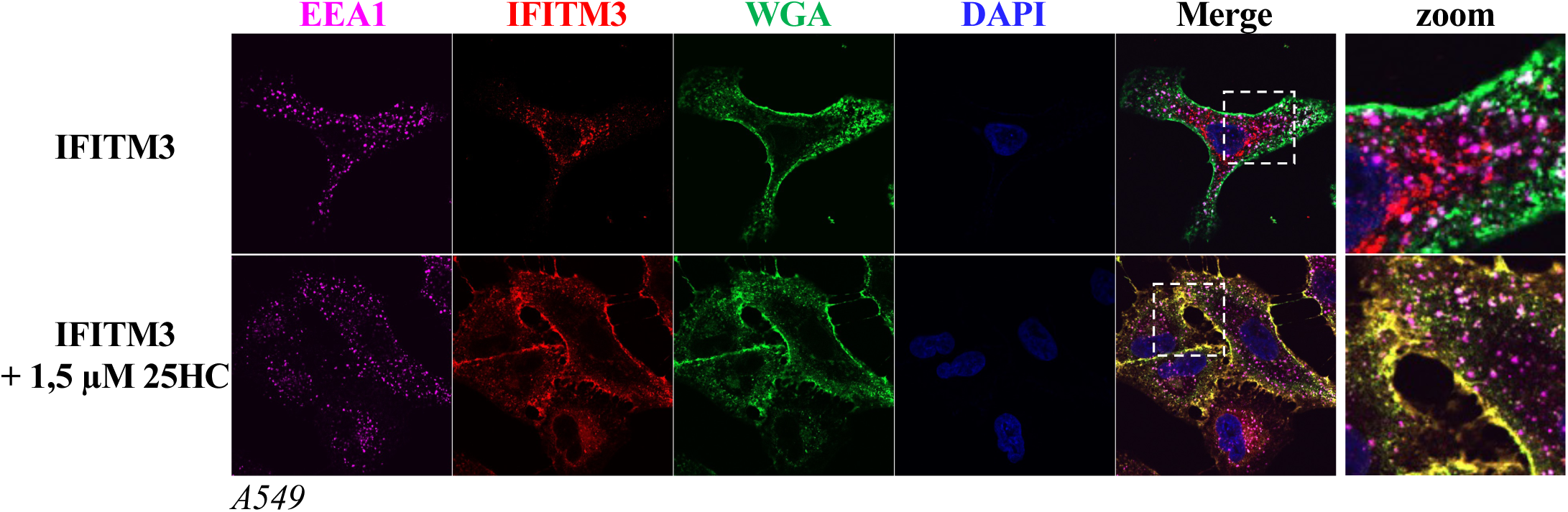
25HC leads to a decreased level of early endosomal recycling of IFITM3 from the plasma membrane. Dox. inducible IFITM3-expressing A549 cells were incubated for twenty-four hours with dox. and/or 25HC prior to confocal microscopy analysis. The picture present typical results obtained.

### The CH25H–25HC axis enhances secretion of IFITM3 in exosome-like vesicles

IFITM3 has been previously detected in exosomal vesicles, in the absence of viral infection (25–27). We therefore tested whether its increased surface localization in the presence of 25HC or CH25h affects its release in extracellular vesicles. To this end, we purified exosomal vesicles from the supernatant of A549 cells expressing IFITM3 alone or in combination with CH25h or from cells expressing IFITM3 and incubated with 25HC. The supernatant was filtered and purified by ultracentrifugation through a 25% sucrose cushion and pellets were then resuspended and analyzed by WB and densitometric quantification. Under these conditions, the levels of vesicle-associated IFITM3 increased when cells were treated with 25HC, and more strongly, upon coexpression of CH25h (3.3 versus 9.2 fold, after normalization with EF1α, Figure 6A and 6B). These findings indicate that CH25h promotes the extracellular release of IFITM3, likely as a downstream consequence of its altered trafficking. To demonstrate that IFITM3 is normally secreted upon association to exosomal vesicles, the supernatant of cells overexpressing IFITM3 was purified by ultracentrifugation through sucrose and then analyzed by immune-gold electron microscopy (Figure 6C). Under these conditions, a readily detectable association of IFITM3 was observed with exosomal vesicles, in agreement with previous data in the literature indicating that, in the absence of viral infection, IFITM3 is a *bona fide* exosomal protein.

**Figure 6.**
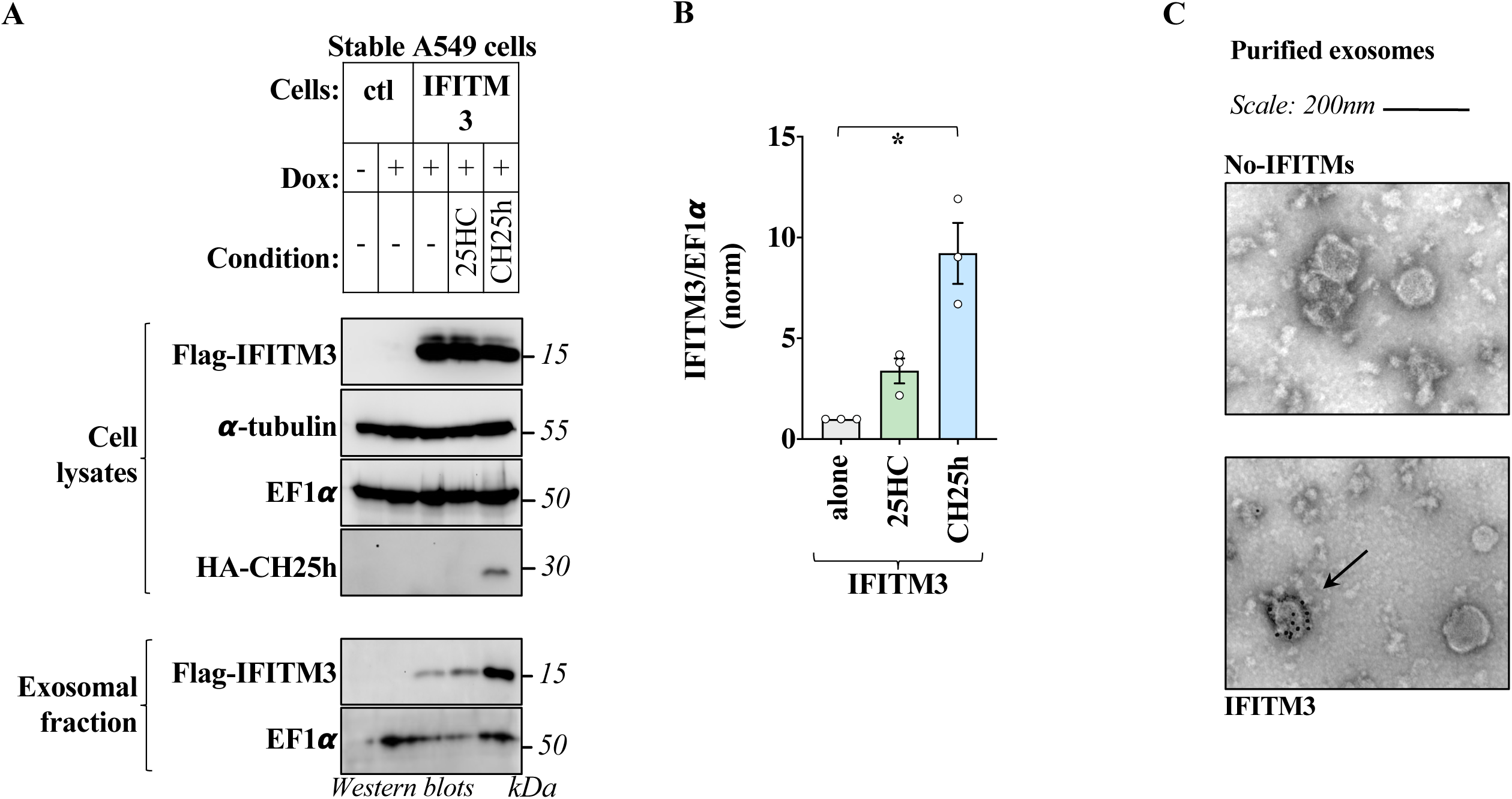
The CH25h-25HC axis leads to an increased accumulation of IFITM3 in exosomal vesicles. Single IFITM3 or double IFITM3/CH25h stable A549 cells were induced for twenty-four hours with dox. prior to harvesting of both cell lysates and supernatants. When indicated, IFITM3 stable cells were either left untreated or co-stimulated with 25HC. After removal of cellular debris by syringe-filtering, exosomal vesicles were purified by ultracentrifugation through a 25% sucrose cushion, prior to WB analysis and densitometric quantification of the protein levels. The WB panels present typical results obtained (A), while the graph presents AVG and SEM of the levels of IFITM3 present in the exosomal fraction, following normalization for EF1α as quantified in 3 independent experiments (B). *, p<0.05, following an Ordinary one-way Anova, Sidak’s multiple comparisons test between the indicated conditions. C) Supernatants produced from control or IFITM3 expressing cells were purified as described before and were then examined by electron microscopy and immuno-gold labeling using an anti-Flag antibody. The panels present representative images obtained of over 400 vesicles scored and the arrows point to typical IFITM3-positive exosomes.

## Discussion

In this study, we examined how two interferon-stimulated membrane-regulating antiviral factors, namely IFITM3 and CH25h, interact functionally during infection with two unrelated enveloped viruses, HIV-1 and VSV. Our analyses reveal that although both restrict viral entry with comparable qualitative outcomes, their antiviral activities are largely redundant. Apparent synergistic effects were observed following spreading infection of HIV-1 in IFITM3-expressing SupT1 cells exposed to 25HC, but in light of the relatively long-time span of the assay, we believe these effects are more likely due to an indirect effect of 25HC on the cell physiology, rather than to antiviral inhibition per se.

Consistent with previous reports, both IFITM3 and CH25h potently inhibit HIV-1 and VSV infection by limiting the entry of virion particles into the cell, which represents the step of the viral life cycle most studied in the literature (4–8, 16, 18, 19, 28). To add to these reports, our results reveal that CH25h mediates a previously-undescribed negative effect during the late phases of the HIV-1 life cycle. Although we have not characterized it in further detail here, this finding posits CH25h as the second antiviral factor capable of inhibiting the viral life cycle of HIV at both early and late phases. It is of interest that IFITM3 acts similarly at both early and late phases of the life cycle of different viruses and that both CH25h and IFITM3 act by altering the behavior of membranes.

In the case of CH25h and 25HC the defect at the late phases of the viral life cycle may be due to an interference with cholesterol-rich virion assembly platforms essential for efficient budding and envelope incorporation (29), or else may be due to an indirect effect on envelope processing as reported for the Lassa virus glycoprotein (30). Interestingly, a connection between lipid metabolism and IFITM3 has been suggested a few years ago, following the report of an interaction between IFITM3 and the VAMP (vesicle-associated membrane protein)-associated protein A (VAPA) (14). VAPA is an important endoplasmic reticulum (ER) adaptor required for the transport of several classes of lipids from the ER to the Golgi (31). The IFITM3-VAPA interaction was reported to interfere with such transport, resulting in a specific increase in the intracellular levels of cholesterol that could have explained the membrane-rigidifying behavior of IFITM3 (14). Despite the fact that this finding remains controversial (32, 33), the connection between CH25h and IFITM3 reopens the interesting possibility that a connection exists between the antiviral effects of IFITM3 and lipid metabolism.

A major and unexpected finding of our study is that the CH25h/25HC axis induces a robust redistribution of IFITM3 from endosomal compartments to the plasma membrane. This relocalization was observed with both exogenous 25HC as well as upon CH25h expression and occurred across multiple cell types. The effect likely reflects a perturbation of IFITM3 re-endocytosis from the plasma membrane, as evidenced by the marked loss of IFITM3 colocalization with early endosomes. Given that IFITM3 undergoes continuous cycles of plasma membrane insertion and internalization, CH25h/25HC-mediated disruption of these steps may alter the turnover of IFITM3 at the membrane and/or modulate its interactions with other cellular factors. These changes do not affect the overall protection conferred by IFITM3 against viral infection. However, they lead to a strong increase in the amount of IFITM3 associated to secreted exosomal vesicles. This increase cannot be explained by differences in the expression levels of IFITM3, which remain steady, and can be more easily explained as a direct consequence of an accrued residence of IFITM3 at the plasma membrane.

While during viral infection we and others have demonstrated that IFITM3 can be readily incorporated into particles of a broad spectrum of viruses (9–11, 28), a few reports have indicated that IFITM3 can also be incorporated in exosomal vesicles in the absence of infection, where they can contribute to exosome-mediated cell-to-cell communications (25, 26). Our findings not only support these previous results, but also extend them by indicating that the CH25h-25HC axis can significantly contribute to drive an increased presence of IFITM3 in this extracellular pool of protein. A functional contribution of exosomal-associated IFITM3 has been reported in the transfer of antiviral mediators from one cell to the other, and more recently also in senescence. A recent study highlighted an important contribution of IFITM3 in the senescence-associated secretory phenotype (SASP) (25), a salient feature with which senescent cells influence their surrounding micro-environment. Given that senescence is associated to an inflammatory transcriptional signature and that CH25h is an integral part of this response, these results suggest the possibility that the CH25h-25HC axis drives at least in part the increased accumulation of IFITM3 in exosomal vesicles, thus contributing to the overall phenotypes associated to SASP and to the particular protein composition of SASP vesicles.

Altogether, our findings highlight the membrane system as an important point of convergent viral inhibition for two broad antiviral factors, IFITM3 and CH25h. These factors do not exert additive effects on viral infection at the cell intrinsic level, under the experimental conditions used. However, in light of the modification in the behavior of IFITM3 that we have described here in the presence of the CH25h/25HC axis, this suggests that these proteins may act in concert in a paracrine manner, potentially influencing cell-to-cell communication and antiviral responses in a broader manner.

Finally, devoting two independent cellular effectors to inhibit a single step in the life cycle of several classes of viruses is likely to offer a competitive advantage to the cell against viral antagonists, as in this case viruses will have to bypass two independent roadblocks rather than a single one.

## Materials and Methods

### Cell culture, DNA constructs, antibodies, densitometry and compounds

Human embryonic kidney and human lung epithelial cells (HEK293T and A549, ATCC cat. CRL-3216 and CCL-185, respectively) were cultured in complete DMEM media with 10% Fetal Calf Serum (Sigma cat. F7524), while the T cell line SupT1 (ATCC cat. CRL-1942) was cultured in complete RPMI-1640 media also supplemented with 10% FCS.

Flag-tagged IFITM3 expressed either from a CMV promoter or in a doxycycline-inducible form in the context of the murine leukemia virus-derived (MLV) retroviral vector have been previously described (11). A codon optimized version of the CH25h was fused at the C terminus to mcherry in the context of an HIV-1-based lentiviral vector (11). A catalytically inactive mutant containing the 4 histidine residues at positions 242, 243, 246, 247 of the catalytic site of CH25h changed to glutamic acid was similarly constructed by gene synthesis (Genewiz). A non-mCherry HA-tagged N terminal version was generated by standard molecular biology in the pRetroX-Tight-Puro vector. pcDNA-D1ER coding an ER-targeted enhanced CFP was a gift from Amy Palmer & Roger Tsien (cat. 36325; Addgene (34).

HIV-1 proviral clone NL4-3 was obtained through the AIDS reagents and repository of the NIH, USA. Single round of infection competent HIV-1-based lentiviral vectors were produced upon co-transfection of four different plasmids: HIV-1 Rev, NL4-3 Env; Gag-Pro-Pol and mini-viral genome coding for a GFP reporter (described in (10), see below for procedure).

Pools of cells expressing in a stable manner IFITM3 and/or CH25h were obtained after transduction with a murine leukemia virus (MLV) retroviral –based gene transduction system pRetroX-Tight system (Clontech), using the following plasmids: Gag-Pro-Pol of MLV (TG5349), the VSV-G envelope; MLV-based genomes coding IFITM3, CH25h or the transcriptional transactivator rtTA (as described in (11), see below for procedure).

The following primary antibodies were used for WB or confocal microscopy, as indicated: mouse monoclonals: anti-α-Tubulin, anti-HA, (Cat. T5168, Cat. H3663; Sigma-Aldrich, respectively), anti-EF1α (Cat. CBP-KK1; Millipore), anti-EEA1 (Cat. MA5-14794; Life Technologies). Rabbit polyclonal antibodies: anti-IFITM3 (Cat. 11714-1-AP; Proteintech), anti-FLAG (Cat. Ab52649; Abcam) and anti-mcherry (Cat. Ab52649; Abcam).

The following secondary antibodies were used for WB: anti-mouse, anti-rabbit IgG-peroxidase conjugated (Sigma, Saint Louis, USA cat. A9044 and AP188P). The following ones were used for confocal microscopy: donkey anti-rabbit IgG conjugated with Alexa Fluor 647 or 555, donkey anti-mouse IgG– Alexa Fluor 488 conjugate and donkey anti-Goat IgG-Alexa Fluor 594 (Life Technologies, Carlsbad, USA, cat. A-21207, or A-32795 or A-32974, A-21202 and A-11058).

Wheat Germ Agglutinin conjugated to FITC (FITC-WGA) was purchased from Santa Cruz Biotechnology (Cat. sc-18822); 25HC was purchased from SIGMA (Cat. H1015). Cell viability in different concentrations of 25HC was evaluated by cell counting upon trypan blue exclusion at different days post treatment.

Following WB, the amount of exosomal-associated IFITM3 was quantified by densitometry analysis using the Chemidoc Imaging System (Bio-Rad), following normalization for the levels of EF1α.

### Production of viruses and infections

Single-round of infection-competent HIV-1 used to study the early phases of HIV-1 infection were produced by calcium phosphate DNA transfection of HEK293T cells (Gag-Pro-Pol packaging construct, mini-viral genome, Rev and the NL4-3 HIV-1 envelope: 8:8:0.5:2 μg each per 10cm dish). Supernatants were collected forty-eight hours afterwards and syringe-filtered to remove cellular debris. Virions were purified through a 25% (w/v) sucrose cushion for 1h30min at 28.000 rpm. The number of infectious viral particles was determined by infecting HeLa cells with virus dilutions and by quantifying the number of green fluorescent protein (GFP)-positive cells obtained two-three days after by flow cytometry. The infectious titer equivalent of non-GFP coding viruses was determined against standards of known infectivity by exo-RT (11).

The early phases of HIV-1 infection were analyzed by challenging target cells expressing IFITM3+/− CH25h at MOI comprised between 0.5 and 1 and by quantifying the number of GFP-positive cells 2-3 days post infection by flow cytometry.

The late phases of the HIV-1 life cycle were analyzed as follows. Virion particles were produced in HEK293T cells co-transfected with plasmids coding the HIV-1 proviral DNA (NL4-3) along with CH25h-mCherry alone or in combination with Flag-IFITM3. After forty-eight hours, the virus was harvested and used to challenge TZM-BL reporter cells. The amount of infectious virion particles was quantified after luciferase analysis on cell lysates using the BrightGlow Lysis Reagent (Promega E2620) and the Tecan Spark Luminometer. Infectious virus yields under various conditions were consistently expressed as fold-change compared with paired viral infection conditions in the absence of IFITM3 or CH25h.

Spreading infections with replication-competent NL4-3 viruses in SupT1 cells were also assessed by intracellular p24 assays and flow cytometry. Cells were collected at the indicated time points, fixed and permeabilized using the FIX & PERM™ Cell Permeabilization Kit (Cat. GAS003, Life Technologies), and stained for intracellular p24 using an RD1-conjugated anti–HIV-1 core antigen antibody (clone KC57; Cat. 6604667, Beckman Coulter) prior toFACS analysis.

Replication-competent GFP-coding VSV-Indiana serotype Rhabdovirus (VSV-GFP) bearing an additional transcription cassette between M and G was used to assess the role of the indicated proteins and conditions on VSV spreading infections. In this case, infectious units were calculated based on infections carried out with limiting dilutions of the virus stock followed by flow cytometry analysis eighteen hours post infection.

Infections were carried out with multiplicities of infection (MOIs) comprised between 0,1 and 1. The percentage of infected cells was determined at eighteen hours post infection by flow cytometry on a FACS Canto II (Becton Dickinson, USA).

### Generation of stable cell lines

Doxycycline-inducible stable cells overexpressing Flag-IFITM3 and/or HA-CH25h were generated using the pRetroX-Tight system (Clontech). Briefly, MLV retroviral vectors are produced by calcium phosphate DNA transfection of HEK293T cells with plasmids coding the MLV Gag-Pro-Pol, the VSVg envelope and two pRetro-X based mini-viral genomes: the first coding the protein of interest (Flag-IFITM3 and/or HA-CH25h) under the control of the dox-inducible promoter, the second coding the transcriptional transactivator rtTA (ratio 8:4:4:4 for a total of 20 μg per 10cm dish). Virion particles in the cell supernatant were directly used for cell transduction followed by selection of cell pools thanks to the Puromycin (Sigma, cat. P8833) and G418 (Sigma, cat. G8168) resistance genes carried by the two constructs. Cell pools were maintained as such to avoid clone-dependent variations.

### HIV-1 entry assay according to the Entry/Uncoating assay based on core-packaged RNA availability and Translation system (EURT)

This assay is based on the direct translation of a Luciferase-bearing HIV-1 genome mimic that can be directly translated in the cell, yielding a precise measure of virus entry into the cell cytoplasm (21). Single round of infection NL4-3 Envelope bearing HIV-1 viruses incorporating this reporter (EU-repRNA) were produced by transient transfection of HEK293T cells, as described above. Infections were carried out for 18 hours in the presence of the capsid destabilizing compound PF74 (10 μg/mL, Sigma, cat. SML0835-5), prior to cell lysis and Luciferase activity quantification (Promega, cat. E2620, according to the manufacturers’ instructions).

### VSV entry (RT-qPCRs)

VSV entry was measured by quantifying the levels of viral RNA in cells 20 min after infection by RT-qPCR in the presence of 100 μg/mL of chycloheximide (CHX, Sigma, cat. 4859) to prevent the translation of viral proteins from positive strand viral RNAs that ignite the full amplification of viral RNA. Prior to cell lysis, cells were washed and treated for 10 minutes at 37°C with trypsin to remove non-internalized virions. Total cellular RNA was extracted according to the manufacturer’s instructions (Macherey-Nagel NucleoSpin RNA cat. 740956) and reverse transcription was performed using random hexamers and oligo(dT) with the SuperScript III reverse transcriptase (Thermo Fisher cat. 18080) also following the manufacturers’ instructions. qPCRs were performed on a StepOne Plus real-time PCR system (Applied Biosystems) using the FastStart universal SYBR green master mix (Roche Diagnostics, cat. 4913914001). Primers amplified the GFP gene inserted into the viral genome: GAACGGCATCAAGGTGAACT and TGCTCAGGTAGTGGTTGTCG.

### Preparation of exosomal fractions

Stably-transduced A549 cells were seeded at equal density and allowed to adhere overnight. The following day, the medium was replaced with or without fresh DMEM containing doxycycline (2 µg/mL) to induce transgene expression, with or without 25-HC at a final concentration of 1.5 µM. After 48 hours of induction, culture supernatants were harvested, clarified by low-speed centrifugation, and layered onto 25% (w/v) sucrose cushions. Samples were ultracentrifuged at 28,000 rpm for 1.5 hour at 4°C in an SW41Ti rotor. The resulting pellets containing an enriched exosomal fraction were resuspended directly in 2× SDS loading buffer for SDS–PAGE and further analyzed by SDS–PAGE gel followed by Western blotting.

### Immuno-gold labeling and electron microscopy

Formvar/carbon-coated nickel grids were deposited on a drop of purified, unfixed exosomal preparations produced in the presence or absence of Flag-tagged IFITM3 for five minutes followed by incubation with an anti-Flag antibody (F7425, Sigma, St-Louis, MO), then by incubation with a 1:30 gold-conjugated (10 nm) goat-anti-Rabbit IgG (Aurion, Wageningen, Netherlands) and fixation in 1% glutaraldehyde. Negative staining was performed using 2% uranyl acetate (Agar Scientific, Stansted, UK) followed by transmission electron microscope analyses (JEOL 1011, Tokyo, Japan).

### Confocal microscopy

Cells were seeded on glass coverslips coated with poly-L-Lysine 0.01% (Sigma cat. P4832) after Lipofectamine 3000-mediated DNA transfection, according to the manufacturer’s instructions (Cat. L3000008, Invitrogen). Twenty-four hours later, cells were fixed with PFA (4%), then permeabilized with 0.1% Triton X-100 in PBS and then incubated with the indicated primary and secondary antibodies. Coverslips were stained with a DAPI-containing solution (ThermoFisher cat. 62248) and mounted using the anti-quenching solution Fluormount G (Southern Biotech). Fluorescent confocal images were acquired on a Zeiss LSM 980 AiryScan confocal microscope. Pictures were analyzed with the ImageJ software.

### Statistical analyses

Two-tail Student t tests or One-way Anova tests were calculated using the Prism 8 software, as specified in the legend to figures.

### Softwares

Confocal microscopy: Fiji software (version 2.0.0), Zen (version 2.3, Zeiss) and Imaris 9.2.0 software (Oxford Instruments Group). WB: Image Lab Touch Software (version 2.0.0.27, Chemidoc Imaging System from Bio-Rad). Flow cytometry: FlowJo (version 10.10, BD).

Statistics and graphs: Graphpad Prism8 (8.4.3, Graphpad software, LLC) and Excel (16.16.3, Microsoft).

## Data availability

All relevant data are within the paper.

## Acknowledgments

Y. S. is the recipient of a post doctoral fellowship from the ANRS|MIE (AO-2022-2) and this work has been supported by a grant from the ANRS|MIE (AO-2022-2) to A.C. We thank Fabrizio Mammano at the Université de Tours France, for sharing the intracellular p24 staining protocol. We acknowledge the contributions of the SFR Biosciences (Université Claude Bernard Lyon 1, CNRS UAR3444, Inserm US8, ENS de Lyon) (LYMIC-PLATIM). A.C. is a researcher of the Centre National de la Recherche Scientifique (CNRS). The funders had no role in study design, data collection and analysis, decision to publish, or preparation of the manuscript.

## Author contributions

Y.S. designed, performed and analyzed experiments; JBG and PR performed and analyzed immunogold EM experiments; A.C. designed the study, acquired funding and analyzed data. Y.S. and A.C. wrote the article.

## Appendix Figures

**Appendix Figure A1. Effects of CH25h and IFITM3 expression on HIV-1 infectivity and effects of 25HC on cell growth**.

The data refers to Fig 1. **A**) Intracellular levels of IFITM3 and (**B**) cell numbers of SupT1 cells incubated with the indicated concentrations of 25HC and analyzed at the indicated days (n=3). C) The graph presents the non-normalized infectivity data of the corresponding graph of Figure 1D.

**Appendix Figure A2. Effects of CH25h and IFITM3 expression on VSV**.

The data refers to Fig 2. **A**) The graph presents the non-normalized infectivity data of the corresponding graph of Fig 2A. **B**) A549 cells expressing dox.-inducible forms of IFITM3 and/or CH25h were obtained upon retroviral-mediated gene transduction. Following the proteins induction with dox (1.5 μg/mL) for twenty-four hours, cells were either analyzed by WB or challenged with VSV-GFP prior to flow cytometry analysis 16-18 hours later. N=3; *, p<0.05 following Ordinary one-way Anova, Dunnett’s multiple comparison test.

**Appendix Figure A3. Effects of 25HC on cell viability and infectivity**.

**A**) Scheme of the approach used here. **B and C**) HEK293T cells were incubated with 25HC for twenty-four hours then either analyzed for viability (cell counting and trypan blue exclusion), or susceptibility to infection upon VSV-GFP challenge and flow cytometry analysis. The graphs present AVG and SEM of 3 independent experiments. *, p<0.05, according to a two-tailed, unpaired Student t test between the two indicated conditions.

