## Supplementary figures and images for "Cholesterol Remodeling by CH25h Rewires IFITM3 Trafficking and Secretion Without Enhancing Antiviral Restriction"

### supplemental figures

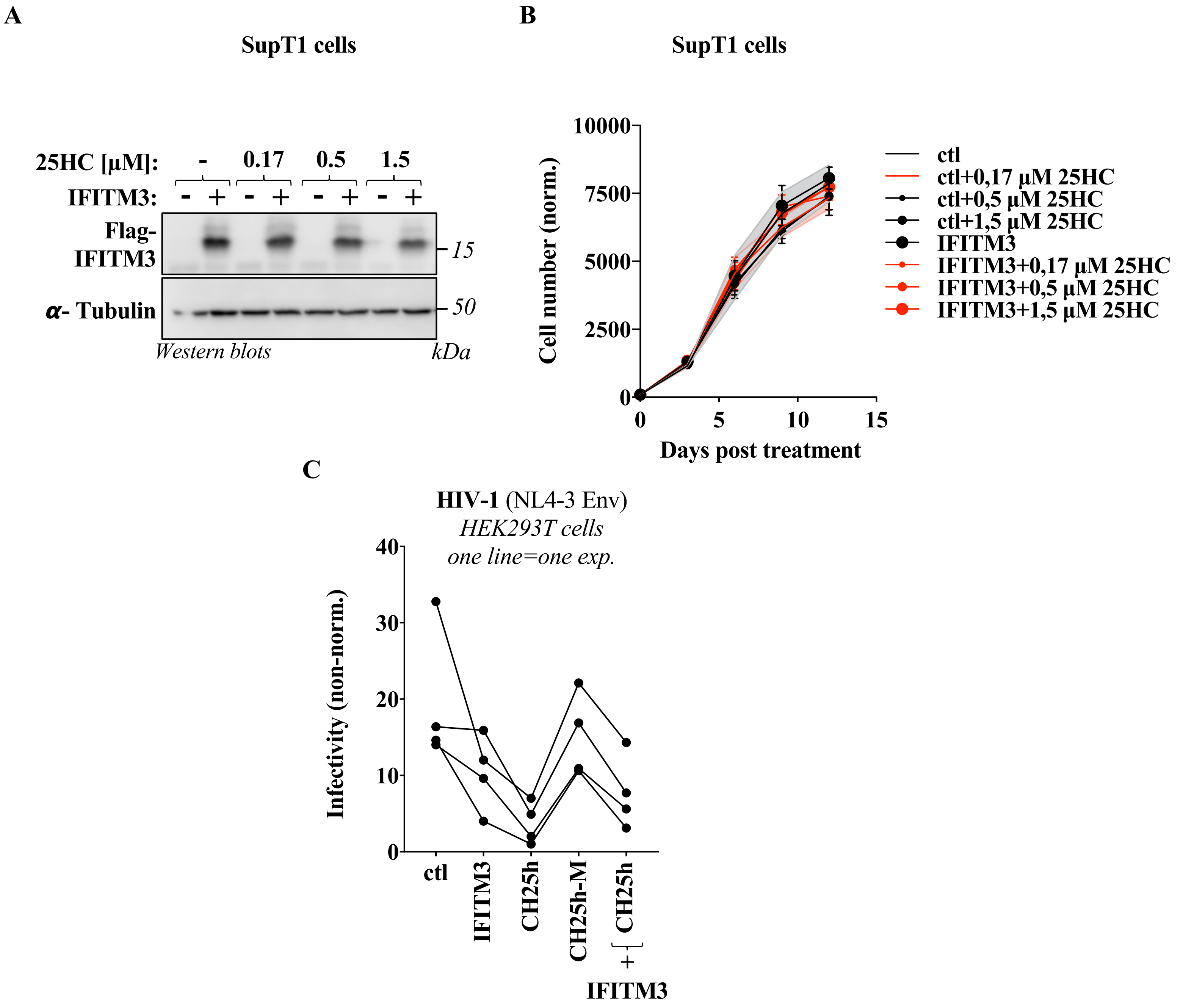

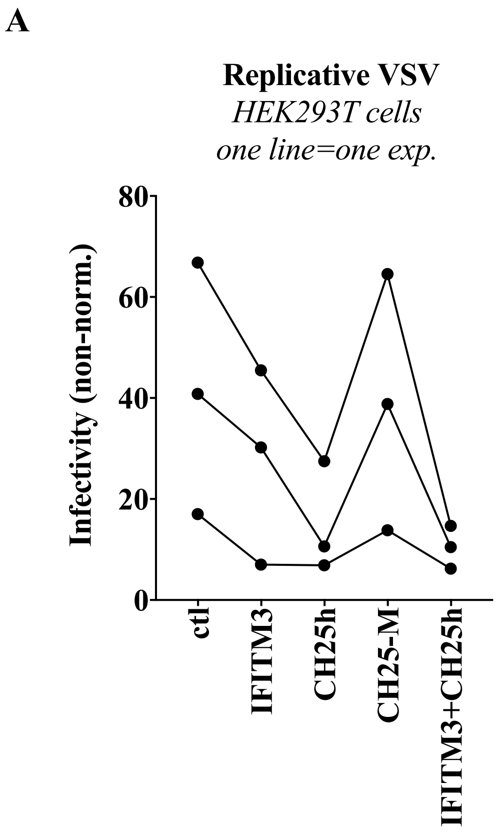

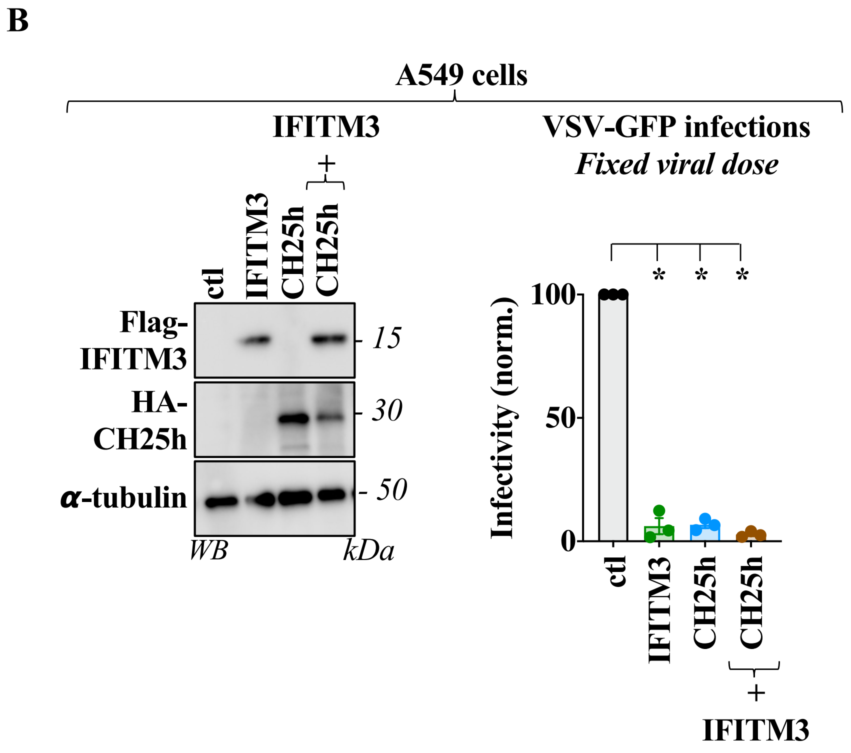

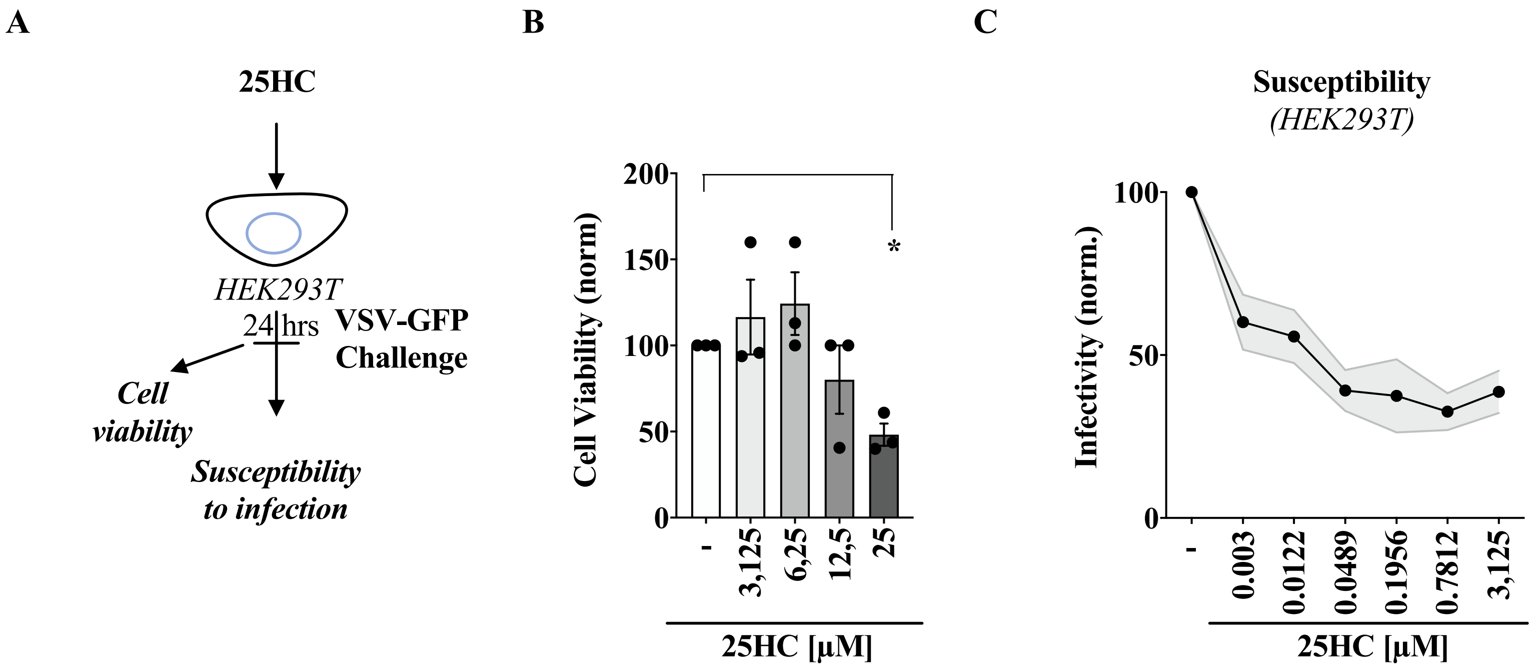
